# OCSANA+: Optimal Control and Simulation of Signaling Networks from Network Analysis

**DOI:** 10.1101/806315

**Authors:** Lauren Marazzi, Andrew Gainer-Dewar, Paola Vera-Licona

## Abstract

**Summary:** OCSANA+ is a Cytoscape app for identifying nodes to drive the system towards a desired long-term behavior, prioritizing combinations of interventions in large scale complex networks, and estimating the effects of node perturbations in signaling networks, all based on the analysis of the network’s structure. OCSANA+ includes an update to OCSANA (optimal combinations of interventions from network analysis) software tool with cutting-edge and rigorously tested algorithms, together with recently-developed structure-based control algorithms for non-linear systems and an algorithm for estimating signal flow. All these algorithms are based on the network’s topology. OCSANA+ is implemented as a Cytoscape app to enable a user interface for running analyses and visualizing results.

**Availability and Implementation:** OCSANA+ app and its tutorial can be downloaded from the Cytoscape App Store or https://veraliconaresearchgroup.github.io/OCSANA-Plus/. The source code and computations are available in https://github.com/VeraLiconaResearchGroup/OCSANA-Plus_SourceCode.

## 1 Introduction

Complex regulatory networks, such as gene regulatory and intracellular signaling networks are ubiquitous in cells. Their computational modeling and analysis is essential to gain understanding of and ultimately, control cellular behavior. Owing to the development of high-throughput measurement technologies and databases, information about signaling network structure is becoming more available, but detailed kinetic parameter information about molecular interactions is still limited, particularly for large-scale complex networks. This lack of knowledge of network dynamics does not preclude researchers from using the topological information of the network to: (1) identify nodes to drive the system towards a desired stable long-term behavior, (2) identify and prioritize combinations of pathway interventions while considering off-target effects, and (3) simulate the long-term dynamic behavior of the system when subjected to perturbations. We previously introduced OCSANA (Vera-Licona et al., 2013), which identifies combinations of pathway interventions. Here, we introduce OCSANA+, an updated Cytoscape app, with the 3 aforementioned analysis capabilities, and an optimized algorithm for the identification of combinations of interventions. To demonstrate the novel capabilities of the tools in OCSANA+ as a pipeline we present two application examples to successfully reproduce and predict simulated and experimental results.

## 2 Features

OCSANA+ (Figure 1) is a user-friendly app for the network analysis and visualization software, Cytoscape (Shannon et al., 2003). It contains 3 main functions:

**Figure 1:**
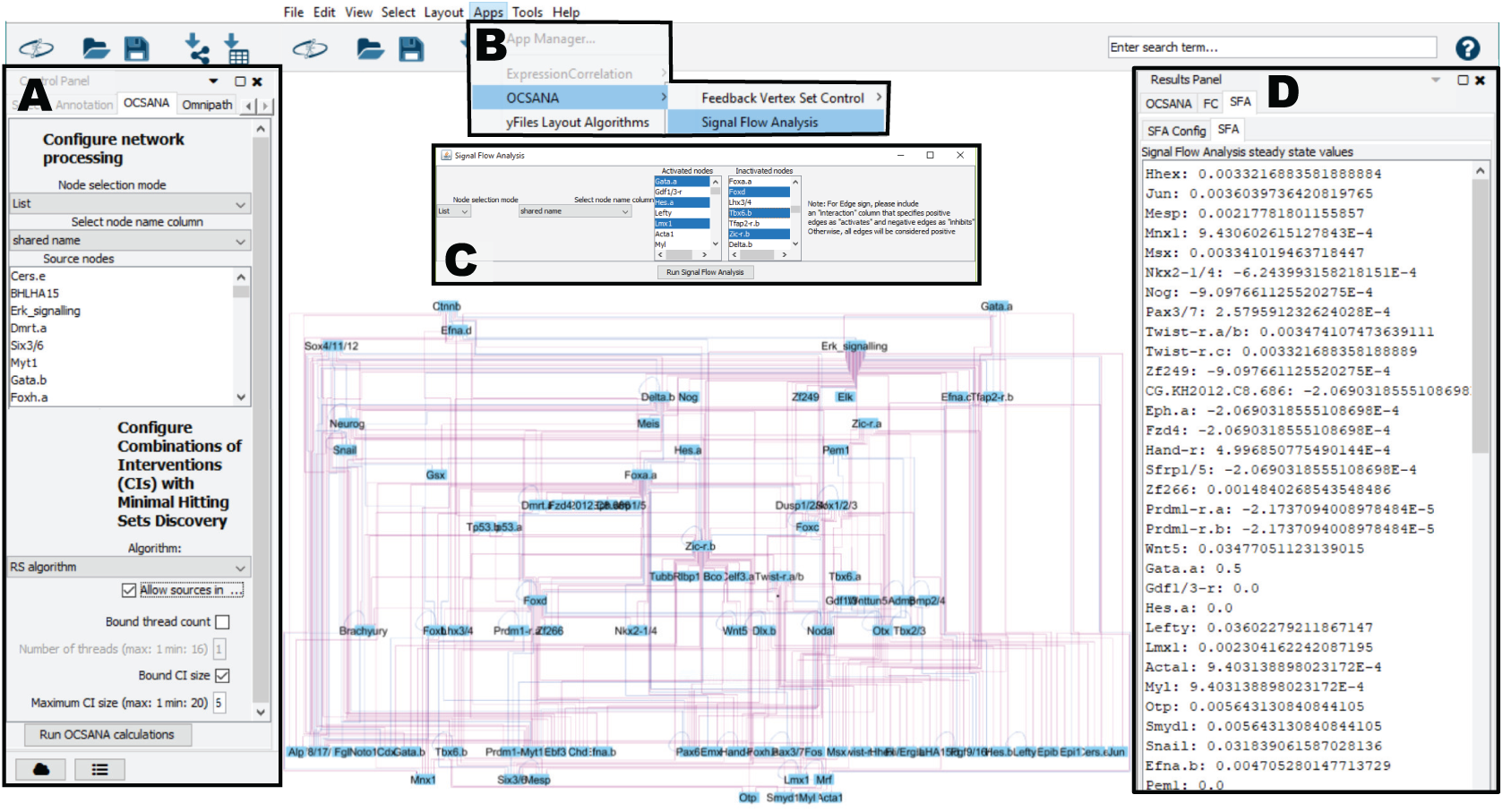
Overview of OCSANA+. (A) OCSANA control interface. (B) Dropdown Selection of FC and SFA algorithms. (C) SFA configuration pop-up menu (D) SFA Results subpanel.

### (1) Combinations of Interventions (CI) Search

A CI is a set of nodes such that each elementary path (a path from user-selected source to target nodes) contains at least one node from this set. This CI set indicates the nodes to be intervened to disrupt all the identified elementary paths. Optimality of CIs is defined in terms of a heuristic scoring. The scoring of a node is based on (i) the lengths of the paths from the node of interest to the targets, (ii) the type of effect on target nodes (e.g. activation/inhibition effect), (iii) side effects with respect to off-target nodes, (iv) the number of elementary paths in which the node participates and (v) the number of targets that such node can reach simultaneously. To identify CIs, users can select from the different algorithms to identify Minimal Hitting Sets (MHSes). In its original version introduced in Vera-Licona et al. (2013), two algorithms to identify MHSes were included: Berge’s algorithm (1989) for an exact solution and a weighted-greedy algorithm for an approximate solution. OCSANA+ includes an additional algorithm, the Reverse Search (RS) algorithm of Murakami and Uno (2014). In Gainer-Dewar and Vera-Licona (2017), the RS algorithm was identified as one of the fastest algorithms from a thorough benchmark of twenty-one algorithms with diverse synthetic and real-world networks. The CI search can be accessed using the OCSANA panel of the Cytoscape control menu.

### (2) Feedback Vertex Set Control (FC)

Structure-based control methods aim to control the system based only on the information provided by the network topology. In OCSANA+ we have included structure-based control methods that apply to systems with non-linear dynamics. Consider a system of *N* nodes, where *i* = 1, …, *N, j* = *N* − *N*_*s*_ + 1, …, *N* and *N*_*s*_ is the number of source nodes (nodes with no incoming edges). At time *t*, the state of the internal node variables *X*_*i*_(*t*) are governed by 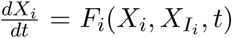 (1) which captures the nonlinear response of node *i* to its predecessor nodes *I*_*i*_,and which includes decay in the dependence of *F*_*i*_ on *X*_*i*_. The state of the source node variables *S*_*j*_(*t*) obeys 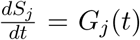 (2). The dynamics described by Eqs. 1 and 2 can be used to describe biochemical dynamics and gene regulation, and are such that they possess some naturally occurring stable states, or dynamical attractors. Dynamical attractors in biological systems like gene regulatory networks can be associated to cell fates or phenotypes. Attractor-based control methods identify control nodes whose manipulation can drive the network from any initial state to any of its attractors. We have implemented two minimal feedback vertex set (FVS) structure-based attractor-based algorithms to control systems with dynamics described by Eq.1 and Eq.2. The minimal FVS (mFVS) of a network is the set of nodes within a graph whose removal would leave a graph without cycles.

i. FC by Mochizuki et al. (2013). For a system described by the nonlinear dynamics of Eq.1, the control action of overriding the state variables of the FVS ensures that the network will asymptotically approach the desired attractor. In this approach, it is assumed that the source nodes of the system converge to a unique state and do not need independent control.
ii. FC by Zañudo et al. (2017). This FC approach extends the method of Mochizuki et al. (2013) to systems in which source nodes are assumed to be governed by Eq.2. As such, in addition to the FVS sets, source nodes of the network need to be steered in the trajectory specified by the attractor.

Using a simulated annealing local search approach (Galinier et al., 2013), OCSANA+ identifies mFVSes in a network. Both FC methods can be selected from the drop-down OCSANA+ menu in the App menu.

### (3) Signal Flow Analysis (SFA)

The SFA algorithm estimates the signal flow (information conveyed by a series of biochemical reactions as represented in a signaling network) based only on the topological information in a signaling network with 60-80% accuracy. Lee and Cho (2018) showed that the ability of the SFA algorithm to predict experimental results was disrupted by randomization of network edges; thus network topology conveys critical information for signal flow, irrespective of kinetic parameters. The algorithm is based on a linear difference equation that considers (i) the activity of a node at the previous time step, (ii) the effect (activating or inhibiting) and influence of incoming edges to a node, and (iii) the initial activities of the node. The user can configure (iii), while (i) and (ii) are calculated from the network topology. Therefore, the estimation of the steady state value for any network node under any perturbation is highly dependent on the correct edge information between network nodes. A user can compare the outputs of different node perturbations to predict the direction of activity change of the network nodes resulting from changes in signal flow. SFA can be selected from the drop-down OCSANA+ menu in the App menu.

## 3 Methods

OCSANA+ is software for the structure-based identification and prioritization of combinations of interventions, attractor-based control nodes and signal-flow estimation of a perturbation on the network. OCSANA+ implements three main algorithms-OCSANA, Feedback Vertex Set Control (FC), and Signal Flow Analysis (SFA). OCSANA+ can be freely accessed and implemented as a Cytoscape App. The OCSANA, FC, and SFA algorithms are fully described in their respective papers (Lee and Cho, 2018; Mochizuki et al., 2013; Vera-Licona et al., 2013; Zañudo et al., 2017). However, we will provide brief descriptions to elucidate their implementation herein.

## 4 Computing Optimal Combinations of Interventions

OCSANA (Optimal Combinations of Interventions from Network Analysis), originally introduced in Vera-Licona et al. (2013), identifies and prioritizes optimal minimal combinations of interventions (CIs) that disrupt the elementary paths from selected source nodes to the specified target nodes. When indicated by the user, OCSANA seeks to additionally minimize the side-effects that CIs can cause on specified off-target nodes. The identification of CIs proceeds as:

1. Pre-processing step: Compute the collection of elementary paths, that is, paths from selected source nodes to selected target nodes. Compute paths from selected source nodes to side-effect nodes, if side-effect nodes have been specified.
2. Score the nodes present in the elementary paths and sort them in a descending order according to OCSANA’s node score. If side-effect nodes were selected, a side-effect score will be incorporated.
3. Compute the so-called minimal hitting sets (MHSs) for the elementary paths according to the selected MHS algorithm and sort them according to OCSANA’s CI score. This sorted list of MHSs is the sought list of prioritized optimal CIs.

### 4.1 Path Analysis

There are two options for identifying elementary paths between source and target nodes.

1. **All non-self Intersecting Paths**: identifies all paths between source and target node that do not include self-loops.
2. **Shortest Paths**: identifies the minimal length paths from source to target for each pair of source and target nodes.

### 4.2 Minimal Hitting Sets

#### Definition 1.1

Given a pair (*V*, Λ) consisting of a universe *V* of *n* elements and a collection Λ of subsets of *V*, a hitting set is a subset *H* of *V* that intersects (hits) every *λ* ∈ Λ. A hitting set *H* is minimal if no proper subset of *H* is a hitting set itself. By mapping the elementary paths (the paths from source nodes to target nodes) onto the Λ collection of subsets of *V*, a hitting set *H* can be seen as a set of nodes such that each elementary path that leads to the target nodes contains at least one node from this set.

Several algorithms have been developed to identify MHSes for a given pair (*V*, Λ). In OCSANA+, we have provided three different algorithms-Berge’s Algorithm and Greedy Heuristic Search (from original OCSANA implementation), and the Reverse Search (RS) algorithm: **Berge’s Algorithm**: Berge’s Algorithm is based in the **Transversal Hypergraph** problem, defined as follows:

Given a finite set *V* of vertices *v*_1_, *v*_2_, … and a set *E* of sets *E*_1_, *E*_2_, …, ⊆ *V* of hyperedges (or “edges”), the pair ℋ = (*V, E*) is a hypergraph. A hypergraph is simple if none of its edges is a subset of any other edge. If *V* = ⋃_*E*∈*ε*_ *E*, as is the case in many applications, we can identify the hypergraph ℋ = (V, E) with the set family E; thus, the theory of hypergraphs is closely related to that of set families.

Readers interested in the full theory of hypergraphs should consult Berge’s monograph on the subject (Berge, 1989).

In the finite hypergraph setting, a (minimal) hitting set of *E* is called a (minimal) transversal of ℋ. The collection of all minimal transversals of ℋ is its transversal hypergraph Trℋ. We note that Tr(Trℋ) = minℋ and, in particular, that Tr(Trℋ) =ℋ in the case that ℋ is simple.

Accordingly, Tr ℋ is sometimes called the dual of ℋ, and the construction of Trℋ is accordingly sometimes called dualization. However, other constructions are also commonly called the hypergraph dual; in particular, it sometimes denotes a hypergraph obtained from H by transforming each edge into a vertex and each vertex to an edge, which coincides with the typical definition of a graph dual. To avoid confusion, we will prefer the term transversal hypergraph throughout.

The core idea of Berge’s algorithm is to proceed inductively over the hyperedges of the hypergraph, alternately adding a new edge to the intermediate hypergraph under consideration and extending the known transversals. We first introduce three important operations on hypergraphs.

#### Definition 1.2

Let ℋ_1_ = (*V*_1_, *ℰ*) and ℋ_2_ = (*V*_2_, *ℰ*_2_) be two hypergraphs. Their join ℋ_1_ ⋁ ℋ_2_ is the hypergraph with vertex set *V* = *V*_1_ ∪ *V*_2_ and edge set *ℰ* = *ℰ*_1_ ∪ *ℰ*_2_. Their meet ℋ_1_ ∧ ℋ_2_ is the hypergraph with vertex set *V* = *V*_1_ ∪ *V*_2_ and edge set *ℰ* = *A* ∪ *B*|*A* ∈ *ℰ*_1_, *B* ∈ *ℰ*_2_.

#### Definition 1.3

Let ℋ be a hypergraph with vertex set *V* and edge set *ℰ*. Its minimization (or simplification), minℋ, is the hypergraph with vertices *V* and edges {*A* ∈ *ℰ*|*ℰ* contains no superset of *A*}. In other words, minℋ retains exactly the inclusion-minimal edges of ℋ. ℋ is simple if ℋ = minℋ. These two operations interact nicely with the transversal construction.

#### Theorem 1

Let ℋ_1_ and ℋ_2_ be hypergraphs. Then the following relations hold:

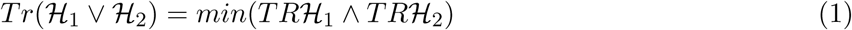

and

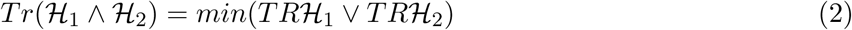

Algorithm Berge then proceeds as follows (Algorithm 1):

1. Let ℋ be a hypergraph with edge set *ℰ* = {*e*_1_, *e*_2_, …, *e*_*n*_} (for an arbitrarily chosen ordering), and for each *i* let ℋ_*i*_ be the subhypergraph of ℋ with all its vertices and its first *i* edges *e*_1_, …, *e*_*i*_.
2. For each *i* in order, compute Trℋ_*i*_ inductively: Trℋ_1_ = {*v*|*v* ∈ *e*_1_} and Trℋ_*i*_ = min(Trℋ_*i*−1_∧Tr *e*_*i*_) = min{*t* ∪ {*v*}|*t* ∈ Trℋ_*i*−1_, *v* ∈ *e*_*i*_} by (1).
3. When the iteration is finished, Tr *H*_*n*_ = Tr*H* by construction.

Berge’s algorithm can be adapted to search only for MHSs of cardinality bounded by some k by simply discarding candidates larger than k.

#### Algorithm 1: Berge’s Algorithm

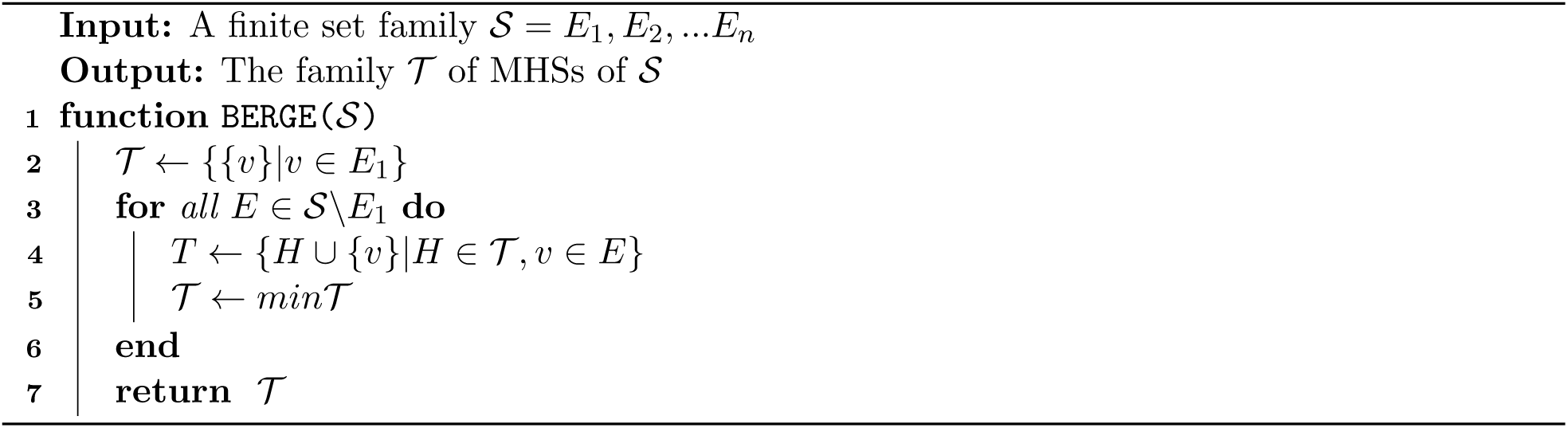

#### Greedy Heuristic

The Greedy Heuristic Approach developed by Vera-Licona et al. (2013) refers to the algorithm designed to compute, according to the selected parameters, the largest possible amount of CIs up to a specified size (or set cardinality). The algorithm starts by building CIs of size 1 and continues building CIs of increasing size until it reaches the specified maximum CI size to be tested. For a given size, the CIs are identified through a weighted-greedy algorithm. The weights of the greedy algorithm are provided by the OCSANA scores of the elementary nodes. The algorithm is initially based on the **MTMiner** algorithm of Hébert et al. (2007). We follow the explanation of the algorithm in Elbassioni et al. (2014) which avoids the algebraic complexity of the original.

Fix a set family 𝒮 = *E*_1_, *E*_2_, …, *E*_*n*_ with underlying element set *V* = ⋃_*E*∈𝒮_, *E* = {*v*_1_, *v*_2_, .., *v*_*m*_}. The MTMiner algorithm is initialized with the set *C*_1_ = {{*v*}|*v* ∈ *V*} of element sets of size 1. At each step of the algorithm, the set *C*_*i*_ of candidate hitting sets of size *i* is processed. First, any set in *C*_*i*_ which is a hitting set is removed and presented to the user as output; as will be seen, it is guaranteed to be minimal. Each remaining set is not a hitting set, so extensions to sets of size *i* + 1 are considered. For each pair (*C, C*_0_) of sets with the property that |*C* ∩ *C*_0_| = *i* − 1, the set *C* ∪ *C*_0_ is constructed. For each of these unions, the algorithm checks whether more sets are hit by *C* ∪ *C*_0_ than by any of its size-i subsets. If so, *C* ∪ *C*_0_ is added to *C*_*i*+1_. The algorithm terminates no later than *i* = *n*, by which time all MHSs have been output.

The algorithm proceeds as follows:

##### Algorithm 2: MTMiner Algorithm

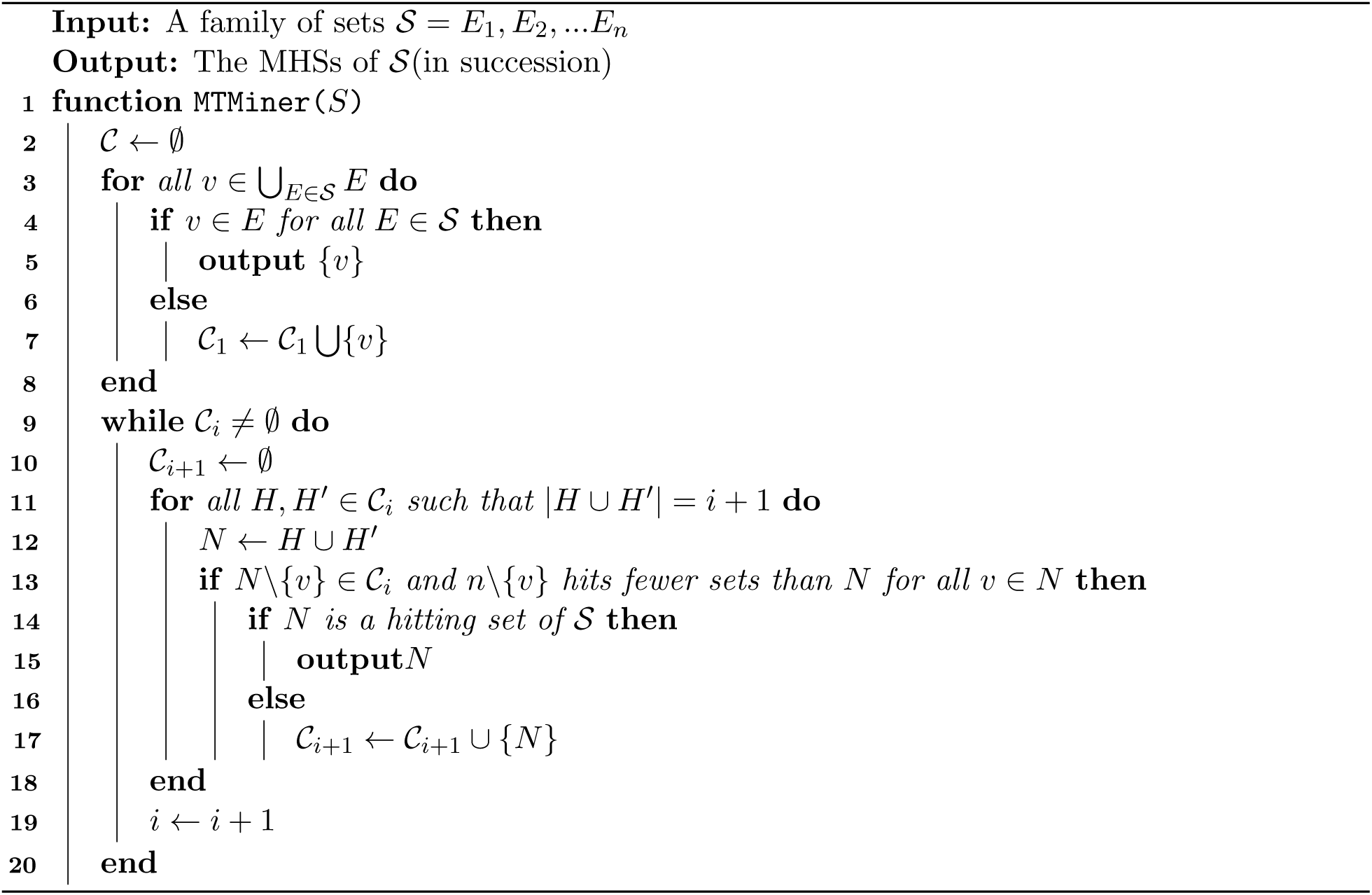

To apply this approach in OCSANA, in lines 13 and 14 we loop over all candidate sets *C* of a given size and consider *C* ∪ {*v*} for every element *V* ∉ *C* such that {*v*} is not itself a hitting set. Additionally, the OCSANA score is used for consideration of sets and elements to optimize the quality of approximate results in cases where complete enumeration is infeasible.

#### Reverse Search (RS) algorithm

In Gainer-Dewar and Vera-Licona (2017), the RS algorithm, developed by Murakami and Uno (2014), was identified as one of the fastest algorithms from a thorough benchmark of twenty-one algorithms with diverse synthetic and real-world networks.

First, we require two definitions. For a given family of sets 𝒮, a sub-MHS is a set which is a subset of some MHS of 𝒮. For a given set *M* of elements of 𝒮 and a given element *m* ∈ *M*, a set *E* ∈ *S* is critical for *m* if *E* ∪ *M* = {*m*}.

##### Theorem 2

If a set *M* is a sub-MHS of a set family 𝒮, then every *E* ∈ *S* is critical for some *m* ∈ *M*. In this case, we say that *M* satisfies the minimality condition.

The RS algorithm, proceeds by incrementally building up sets that satisfy the minimality condition, discarding those which can be shown not to be sub-MHSes and yielding those which become hitting sets.

Let *F* be a hyperedge in the hypergraph ℱ. Let ℱ(*v*) be the set of hyperedges in ℱ that includes *v*. The **RS algorithm** proceeds as follows:

##### Algorithm 3: RS Algorithm

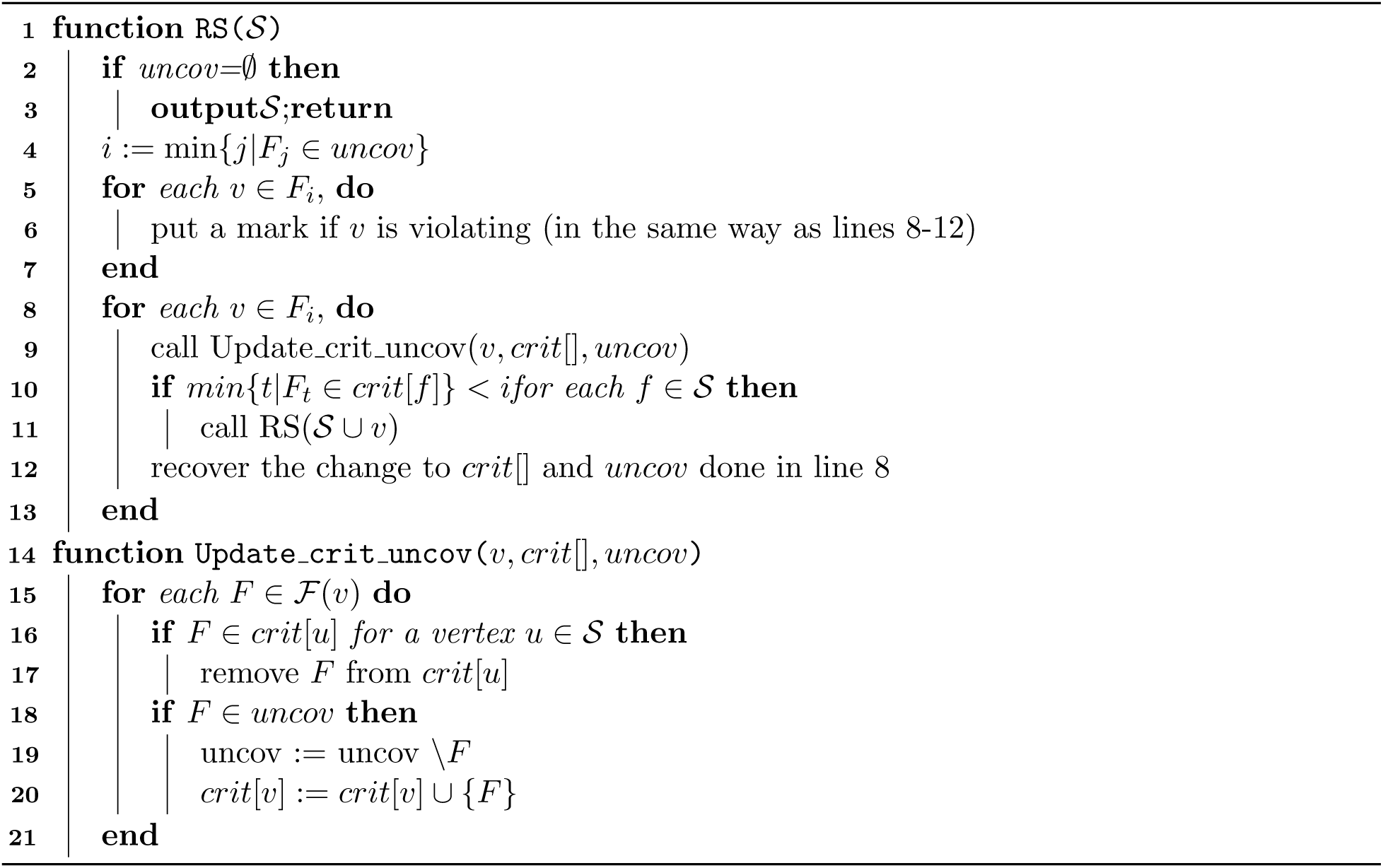

### 4.3 Scoring Elementary Nodes and Combinations of Interventions

The scoring of a node is based on (i) the length of the paths from the node of interest to the targets (ii) the type of effect on target nodes (e.g. activation/inhibition effect) (iii) side effects with respect to off-target nodes (iv) the number of elementary paths in which the node participates (v) the number of targets that such node can reach simultaneously. Details on the OCSANA score can be found in Vera-Licona et al. (2013) Supp1_AlgDescription.

## 5 Computing Structure-Based Attractor-Based Control Nodes with Feedback Vertex Set Control (FC)

Structure-based methods study the controllability of systems based solely on the structure of the network (Ching-Tai Lin, 1974; Mochizuki et al., 2013; Zañudo et al., 2017). In the recent years, structure-based methods for systems with non-linear dynamics have been proposed. One of such structure-based methods for non-linear dynamics is the feedback vertex set control (FC) introduced by Mochizuki et al. (2013). FC is a structure-based control method focused on the controllability of the system by restricting the target states to attractors. Mochizuki et al. (2013) mathematically proved that, for a network governed by non-linear dynamics like those of cell signaling, the control action of overriding the state variables of the feedback vertex set (FVS) into the trajectory specified by a given dynamical attractor ensures that the system will asymptotically approach the desired dynamical attractor.

Consider a directed graph *G* = (*V, E*) comprised of node set *V* and edge set *E*. The node states of *G* are described by the ODE

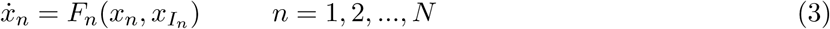

where for the dynamics *x* of node *n* ∈ *V, I*_*n*_ is the set of nodes that regulate node *n*, such that self regulatory loops (*n* ∈ *I*_*n*_) are only positive. Additionally, we assume *F*_*n*_ satisfies decay condition:

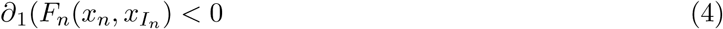

for all n where *∂*_1_ is the partial derivative w.r.t. the first occurrence of *x*_*n*_ and not 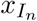.

### Definition 2.1

In *G*, a subset *I* ⊆ *V* of nodes is Feedback Vertex Set (FVS) if and only if a removal of set *G \ I* leaves a graph without directed cycles.

### Definition 2.2

In a dynamic system, a subset *J* ⊆ *V* of nodes is a set of determining nodes if and only if two solutions satisfy 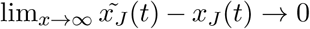 whenever 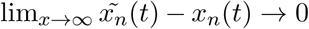 for all components *n* ∈ *J* ⊆ *V*.

In Fiedler et al. (2013); Mochizuki et al. (2013) these two definitions were proved to be equivalent for dynamics in a network. Therefore, observation of the long-term dynamics of the FVS is sufficient to identify all possible attractors of an entire system. Controlling the dynamics of the FVS 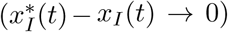 is sufficient to drive the dynamics *x*(*t*) of a whole system to converge on one of any attractors *x*^∗^(*t*).

Zañudo et al. (2017) expanded the FC framework to include graph theory source nodes (nodes with no incoming edges). Whereas Mochizuki assumes that source nodes converge to a unique trajectory and do not need independent control, Zañudo asserts that source nodes can denote external stimuli a system may respond to, which could result in different attractors for source node states. Under this framework, assume the dynamics of directed graph G are governed by Eq. 3 for internal nodes((*N* − *N*_*s*_) where *N*_*s*_ is the number of source nodes in the network), and

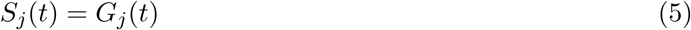

where *S* ⊆ *V* is the set of source nodes, and the dynamics of each source node *j* is independent of the internal node variables, fully determined by *G*_*j*_(*t*), and does not include a decay term.

In OCSANA we have included two structure-based FVS-based attractor-based methods for non-linear systems:

1. FC control from Mochizuki et al. (2013). In this type of control it is assumed that the system does not have source nodes or that source nodes converge to a unique state and do not need an independent control.
2. FC control from Zañudo et al. (2017). This control approach is an adaptation of Mochizuki’s FC control: The control of the source nodes and of the FVS of a network is needed to guarantee that the system can be driven from any initial state to any of its dynamical attractors. The addition of the control of source nodes is under the assumption that the state of the source nodes can affect the dynamical attractors available to the system.

### 5.1 Identifying the Minimal Feedback Vertex Set

The minimal Feedback Vertex Set problem is a well known NP-hard problem. Many algorithms have been developed to find the near-minimum FVS. Based on the implementation of FC in Zañudo et al. (2017), we have used a simulated annealing local search approach, **SA-FVSP**, originally described in Galinier et al. (2013). Local search algorithms, when applied to combinatorial optimization problems define a search space (the set of configurations), an evaluation function, and a neighborhood function. To perform local search for FVSes, the set of configurations is any ordered sequence S of vertices such that, if two endpoints of an arc belong to the sequence, the starting point appears earlier than the endpoint (Algorithm 4). The FVS is *G \ S*. A simulated annealing algorithm is used to explore the neighborhood of S. SA-FVSP has been show to outperform the greedy adaptive search procedure by Pardalos et al. (1998).

#### Algorithm 4: SA-FVSP Algorithm

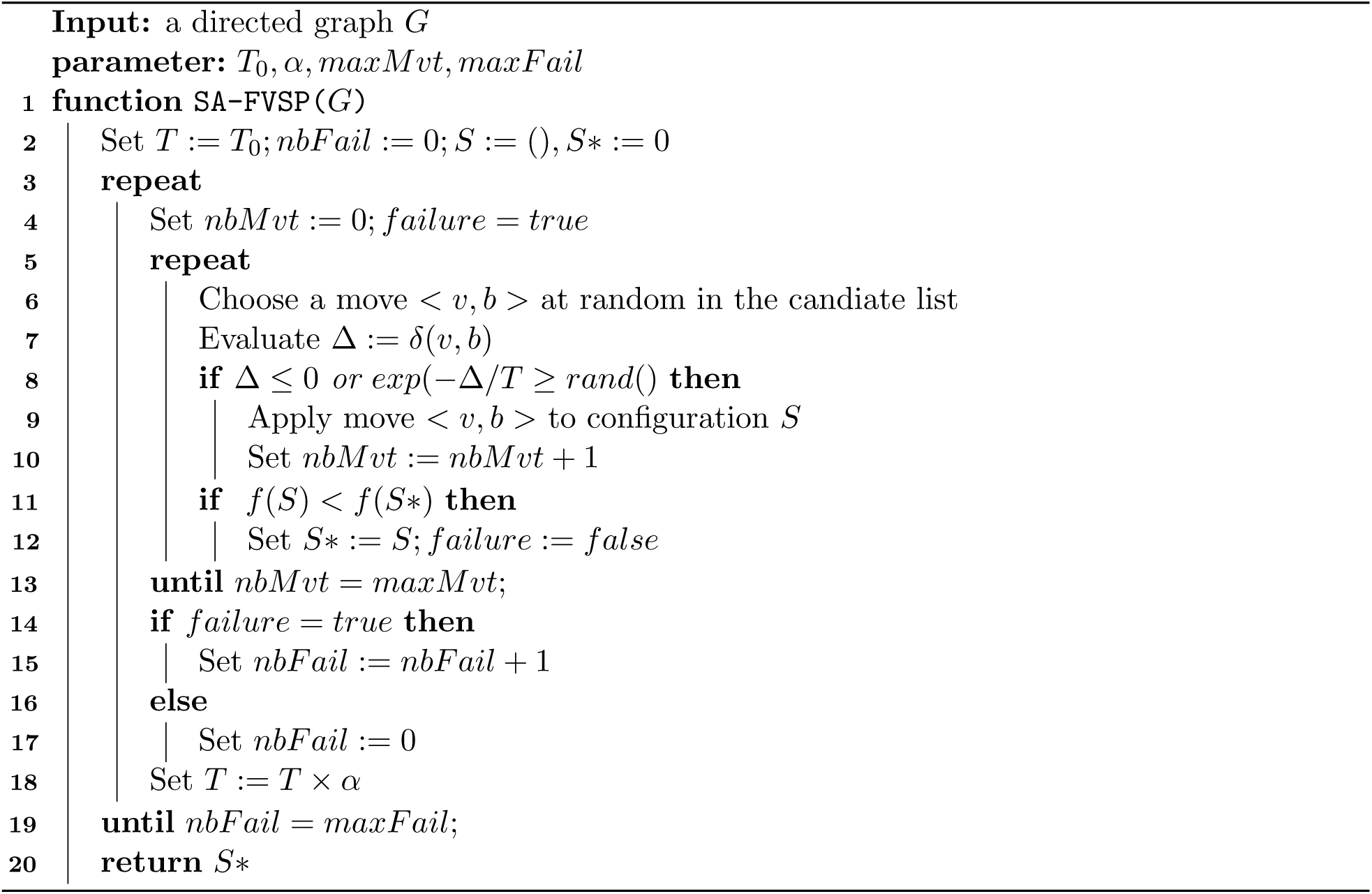

Our Java implementation of the Python code for FC provided by Zañudo et al. (2017) can be found in the OCSANA+ github repository (https://github.com/VeraLiconaResearchGroup/OCSANA-Plus_SourceCode).

## 6 Estimating Signaling Network Perturbations’ Effect with Signal Flow Analysis (SFA)

The SFA algorithm estimates the signal flow (the information conveyed by a series of biochemical reactions as represented in a signaling network) based only on the topological information in a signaling network. Signal Flow Analysis is based on the signal Propagation algorithm, described in Lee and Cho (2018).

The activity *a* of a node *i* is determined by the activities of its regulators and the basal activity of the node. This can be defined as

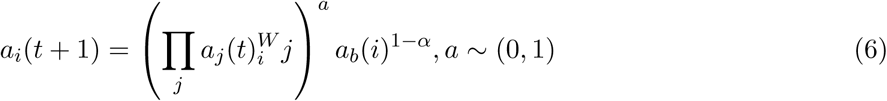

where *a*_*i*_(*t*) and *a*_*b*_(*i*) ∈ *P* are the activity at time *t* and the basal activity of node *i, W*_*ij*_ ∈ *P* is the weight of an edge between node *j* and node *i*, which represents how much node *j* and node *i* through the edge, and *α* ∈ *P* is a hyperparameter for weighted multiplication (default=0.5). The effect of input stimulation is reflected to the basal activity of input nodes. When taking the

logarithm of Eq.6, it becomes a linear difference equation

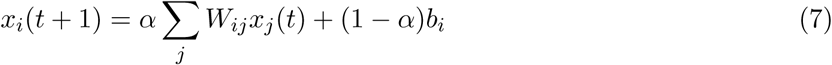

where *x* = *log*(*a*) ∈ *P*^*N*^, *b* = *log*(*a*_*b*_) ∈ *P*^*N*^ and *W* = *P*^*N*×*N*^ is the link weight matrix. This is solved exactly for the steady state values (*x*_*s*_) by

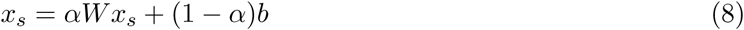

To determine the effect of multiple perturbations the difference between the log steady state values can be calculated as

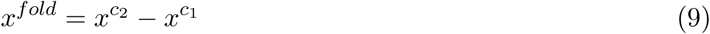

where *c*_*i*_ denotes each condition. Since *x* is the logarithm of the activity at steady state, Eq.9 is essentially a log fold change. The sign (positive or negative) of *x*^*fold*^ denotes if the activity is up or downregulated in the compared conditions, rather than predicting the accurate amount of change.

The OCSANA+ Java implementation of the Python code of Lee and Cho (2018) can be found in the OCSANA+ github repository (https://github.com/VeraLiconaResearchGroup/OCSANA-Plus_SourceCode/tree/master/OCSANA_Plus_SourceCode).

## 7 Results: Application Examples

We demonstrate the ability of OCSANA+ to successfully reproduce simulated and experimental results from two biological signaling networks with non-linear dynamics. Furthermore, we show how OCSANA+ can improve the simulated results. All computations are available in https://github.com/VeraLiconaResearchGroup/OCSANA-Plus_SourceCode.

## 8 Application Example 1: Ascidian embryo cell fate specification

To show experimentally the ability of FC nodes to control cell fates, Kobayashi et al. (2018) identified FC nodes in the gene regulatory network of *Ciona intestinalis* embryos and performed *in vitro* knock-down and upregulation experiments of the FC nodes to control cell fate specification.

### 8.1 Description of the network and FC Analysis Performed in Kobayashi et al.(2018)

The Gene Regulatory Network (GRN) for the specification of cell fate was determined by a genome-wide gene knockdown assay for regulatory genes that are expressed during embryogenesis (Imai et al., 2006) and that was recently updated using data that had been accumulated after the initial construction (Satou and Imai, 2015). This GRN contained 92 genes (nodes) and 328 regulatory interactions (edges) (Figure 1).

They identified all 12 of the minimal Feedback Vertex Sets (FVS) found in the network, each comprised of 5 genes. After selection of 1 FVS containing nodes *Foxa.a, Foxd, Neurog, Zic-r.b*, and *Erk signaling*, morpholino antisense oligonucleotides and synthetic mRNAs were used to knock-down and up-regulate gene activity, respectively. RT-qPCR was performed for all combinations of perturbations (2^5^ = 32). 22 of the perturbations were then classified into their predominant cell fate (epidermis, brain+pan-neural, pan-neural, endoderm, notochord, mesenchyme) based on marker expression patterns as compared to an unperturbed control embryo (Table 1). For each tissue fate, one perturbation was selected for *in-situ* hybridization experiments.

**Table 1:**
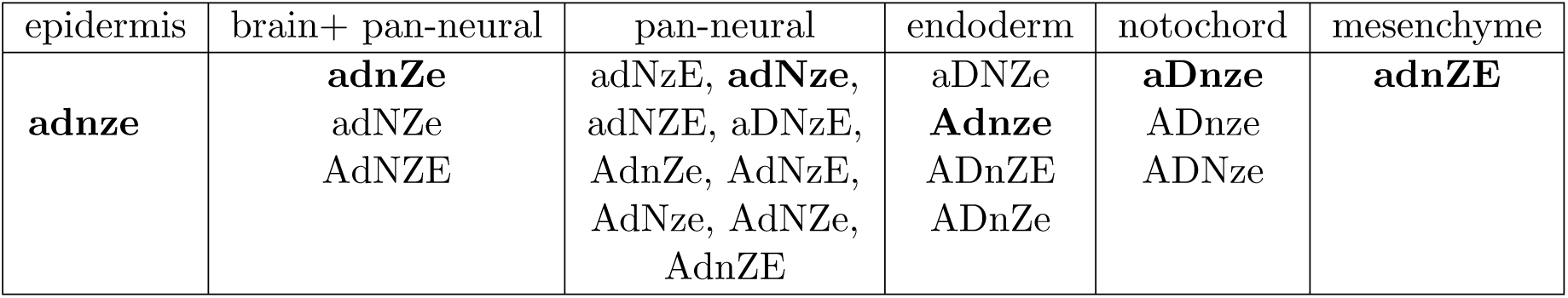
Marker expression of experimental embryos. Each experimental condition is represented by a five-letter code in which up-and down-regulation of *Foxa.a, Foxd, Neurog, Zic-r.b, and Erk signaling* are represented by A/a, D/d, N/n, Z/z, and E/e, respectively. Perturbations in bold letters were selected for *in-situ* hybridization experiments. Reproduced from Kobayashi et al. (2018)

### 8.2 Analysis on the Ascidian Embryo Cell Fate Network using OCSANA+

After performing the *FC without source nodes* analysis through OCSANA+ we identified the same 12 minimal FVSes as in Kobayashi et al. (2018). To reproduce the RT-qPCR and *in-situ* hybridization results in Kobayashi et al. (2018), we used SFA to predict the effect of perturbations (up- or downregulation) of the 5 FVS nodes. We additionally simulated an unperturbed state using initial values of activated *Gata.a* and *Zic-r.a* which are noted to initiate the zygotic developmental program (Kobayashi et al., 2018, Satou and Imai, 2015). Since the values produced by SFA are qualitative and not quantitative, we can calculate the logFC as the difference between perturbed and unperturbed log steady state values which is equivalent to the log of the ratio of the perturbed and unperturbed steady state values. The logFC values that are positive indicate up-regulation in the perturbation attractor, while negative values indicate down-regulation in the perturbation attractor. Our simulations were deemed successful if the perturbation logFCs was upregulated for the correct marker gene for the specified tissue (Table 2). For four of the six perturbations (66%), we were able to predict the correct upregulated gene for tissue fate specification.

**Table 2:**
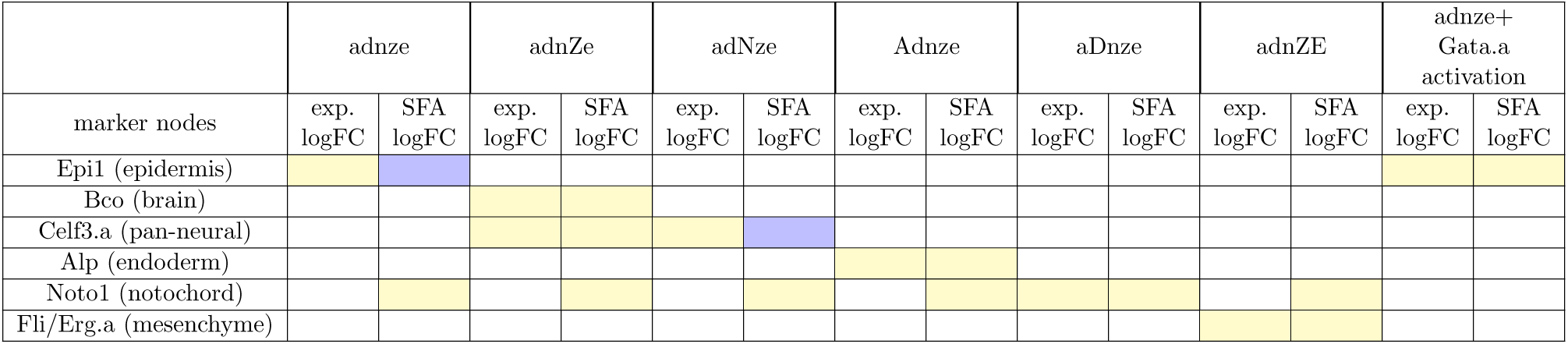
Summary of results of *in silico* simulations of FVS node perturbations. Yellow boxes indicate upregulation of the cell fate tissue marker. Blue boxes indicate downregulation of cell fate tissue marker. For each perturbation, the experimental logFC is compared with the SFA logFC between the perturbed and unperturbed state (perturbed-unperturbed).

Considering that the GRN studied contains source nodes, which in principle can affect the dynamical attractors in the system, we applied the extended FC approach from Zañudo et al. (2017). We identified nine source nodes: *Ctnnb, Gata.a, Gdf1/3-r, Hes.a, Pem1, Sox4/11/12, Tp53.a, Tp53.b, Zic-r.a*. In addition to being critical for zygotic development, *Gata.a* and *Zic-r.a* are required for the specification of ectodermal tissue and mesenchyme tissue, respectively (Satou and Imai, 2015).

We used OCSANA in OCSANA+ to canalize the signal from the network source nodes to a specified cell fate. For example, if we want to predict additional nodes that may control the signal to epidermal specification, we identify combinations of interventions that can be used to intervene in paths from the nine source nodes to epidermis marker *Epi1*. Configuring path finding for *all non-self-intersecting paths* with a maximum length of 20 nodes from source to target, and CI discovery with the RS algorithm, allowing source nodes to be in CIs, we identify a CI of two nodes: source node *Gata.a*, and FVS node *Erk signaling*. SFA was used to predict the effect of downregulation of all FVS nodes perturbation (adnze), and activation of *Gata.a* (Table 2). The addition of *Gata.a* activation then produced a steady state value for epithelial marker *Epi1* that matched the experimental result. However, neither addition of the source nodes, nor a CI from source nodes to pan-neural marker *Celf3.a* was able to achieve SFA simulation that upregulated *Celf3.a*.

## 9 Application Example 2: Drosophila Segment Polarity Genes Network

To assess the extended FC approach with source nodes, Zañudo et al. (2017) used the *Drosophila Melanogaster* segmentation polarity genes network that guide gene expression during embryonic development. The purpose of this study was to show the success of FC to steer the dynamics of a system towards any natural attractor in a validated model. With an established Boolean Model of the Segment Polarity gene network, the authors showed that fixing the values of the FC nodes to their state in the correctly patterned attractor (the wild-type attractor) was sufficient to guide the system to the patterned attractor irrespective of the initial state of all other network nodes.

### 9.1 Description of the segment polarity gene network and FC Analysis Performed in Zañudo et al. (2017)

The fruit fly *Drosophila Melanogaster* segment polarity gene network is a well studied model of embryonic patterning. The fruit fly body is composed of segments, and the segment polarity genes play an essential role in the correct location of appendages, and maintaining boundaries between embryonic sections (Hooper and Scott, 1992). Albert and Othmer (2003) showed that a model solely of the network topology of the segment polarity genes was sufficient to reproduce the dynamics of the segment polarity gene networks. The model constructed by these authors and used by Zañudo et al. (2017) represents four cells as a repeating unit to reproduce wild-type stable cell patterning (Figure 2). This model contains 56 nodes and 144 edges representing the four cells as hexagons with two cell-to-cell boundaries. Zañudo and coauthors focused on the expression of *hedgehog* in cell 2 (*hh*_2_) and *engrailed* in cell 2 (*eg*_2_) at the steady states, as they are the major determinants of embryonic patterning and development.

**Figure 2:**
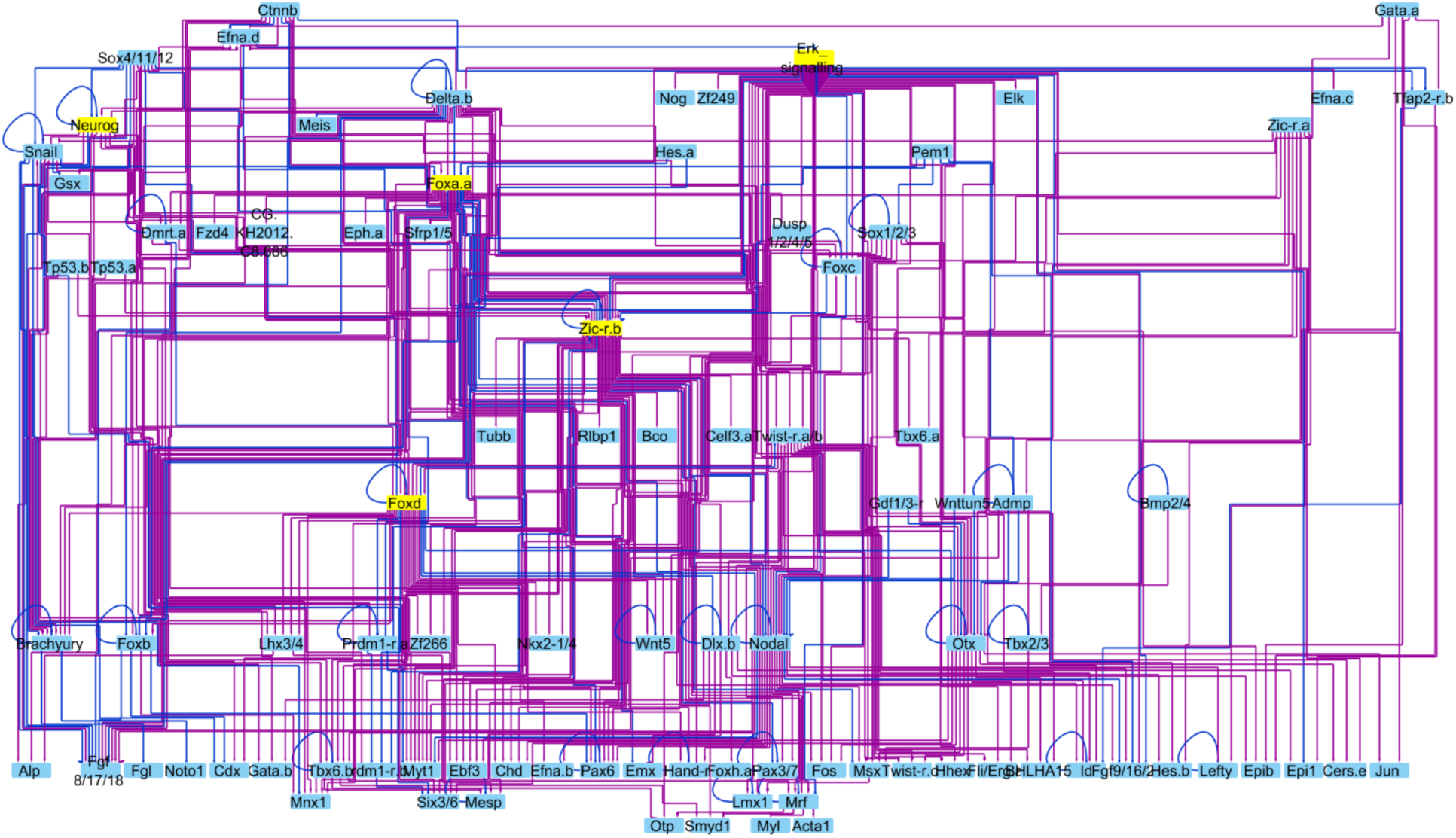
Gene Regulatory Network for Cell Specification in Ascidian *Ciona intestinalis* used in Kobayashi et al. (2018). Purple edges indicate activating interactions, while blue edges indicate inhibitory interactions. Nodes highlighted in yellow are those who are members of the FC set explored in Kobayashi et al. (2018).

Zañudo and coauthors identified two initial conditions for network nodes that yielded either the wild-type patterned steady state, or the unpatterned steady state. They identified an FC consisting of the following nodes: *SLP*_0_, *SLP*_1_, *SLP*_2_, *SLP*_3_, *wg*_0_, *wg*_1_,*wg*_2_, *wg*_3_, *PTC*_0_, *PTC*_1_, *PTC*_2_, *PTC*_3_, *CIR*_0_, *CIR*_2_.

**Figure 3:**
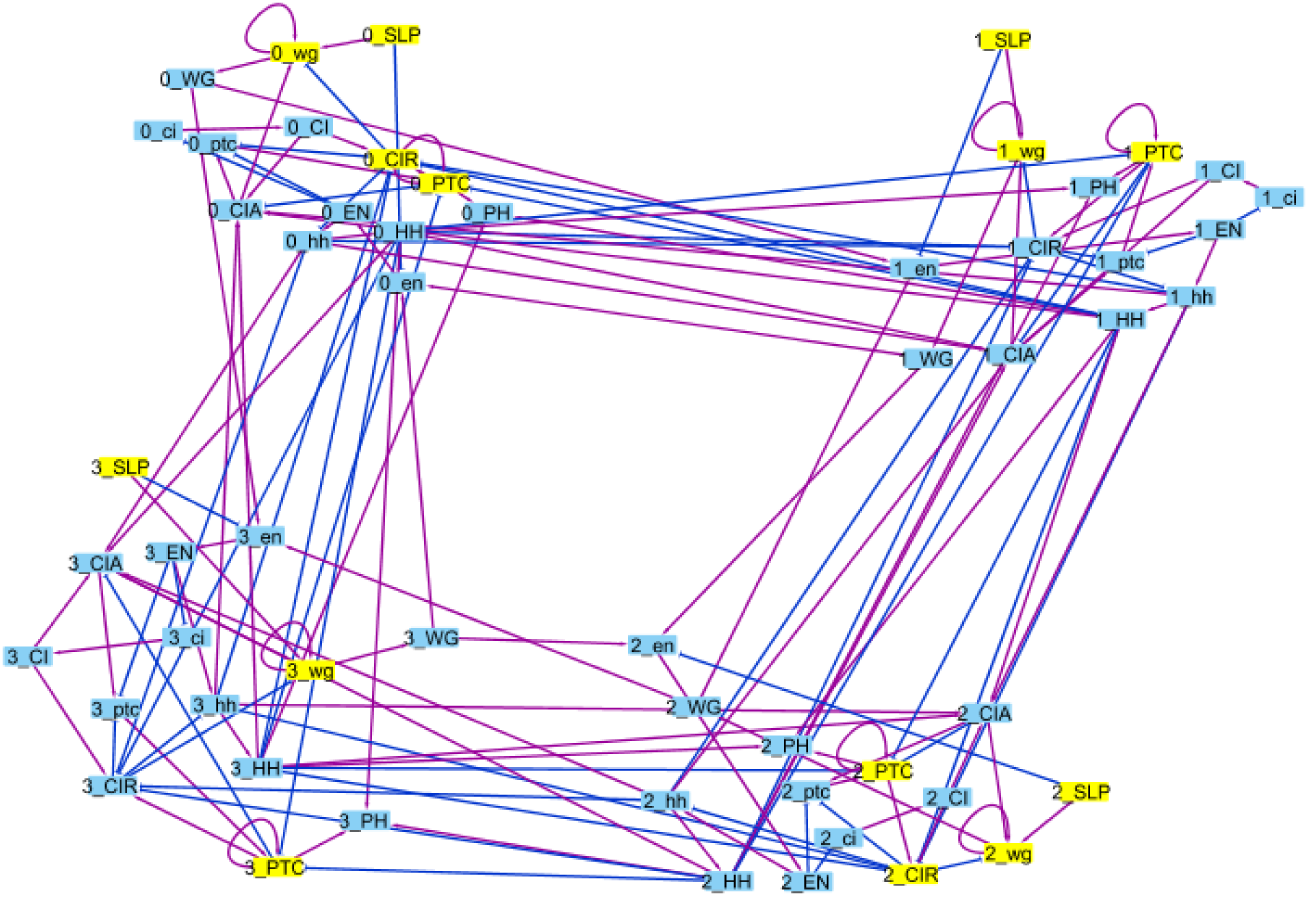
*Drosophila* Segment Polarity network Purple edges indicate activating interactions, while blue edges indicate inhibitory interactions. Nodes highlighted in yellow are those who are members of the FC set explored in Zañudo et al. (2017).

Using a Boolean model derived from the network, Zañudo and coauthors simulated the results of FC perturbations. From any arbitrary initial condition, including the unpatterned initial state, fixing the state of the FC nodes to their values at the wild-type steady state resulted in the wild-type steady state (Table 3).

**Table 3:**
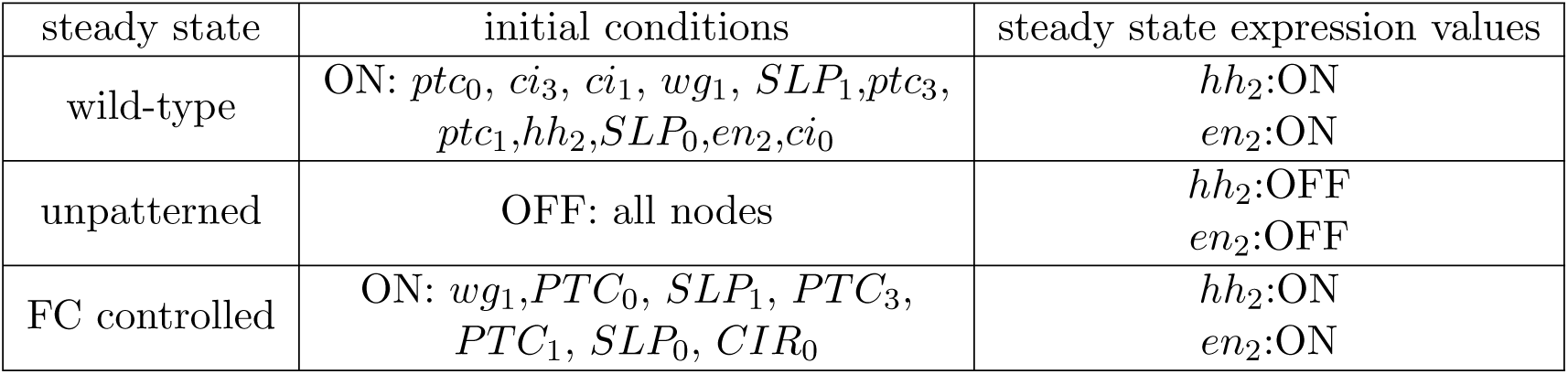
Summary of the results of the Boolean simulations of *Drosophila* segment polarity gene network reproduced from Zañudo et al. (2017).

### 9.2 Analysis on the Segment Polarity Genes Network using OCSANA+

We loaded the static Segment Polarity Network used in Zañudo et al. (2017) into Cytoscape. Using the *FC with source nodes* algorithm through OCSANA+ we identified 6 FCs including the above FC set used for control in Zañudo et al. (2017). To reproduce the control results we simulated the wild-type steady state and unpatterned steady state using SFA through OCSANA+. We calculated the logFC between the wild-type and unpatterned log steady state values for *en*_2_ and *hh*_2_ generated by SFA. Then, we simulated the FC control in the same manner as in Zañudo et al. (2017) by fixing the state of the FC nodes to their values at the wild-type steady state, and leaving all other nodes at the unpatterned state initial values. The logFC between the FC controlled steady state and the unpatterned steady state indicates upregulation of *en*_2_ and *hh*_2_ (Table 4).

**Table 4:**
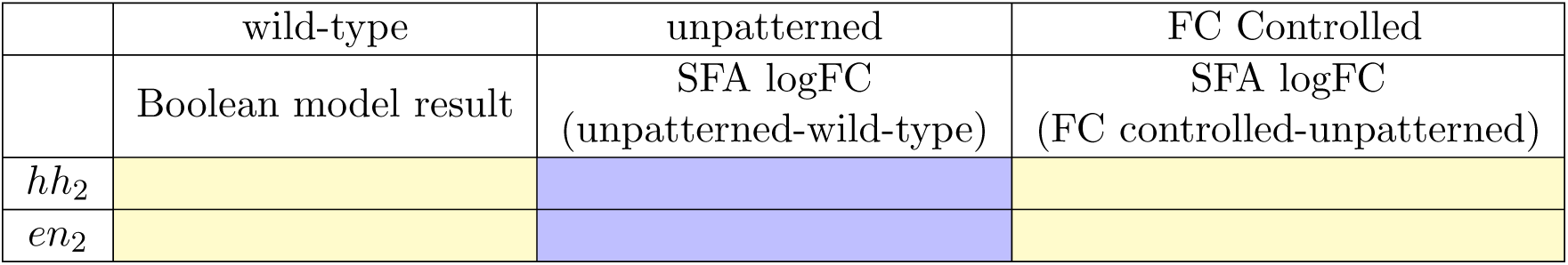
Summary of results of in silico simulations of FC node perturbations. The wild-type column represents the state of nodes in the wild-type attractor (ON). Yellow boxes indicate upregulation of the segment polarity gene. Blue boxes indicate downregulation of the segment polarity gene, as computed from the SFA logFC.

## 10 Discussion

We show that OCSANA+ can be used to accurately predict up to 85% of the experimental results of Kobayashi et al. (2018). We were able to correctly predict the upregulated gene for tissue fate specification for 66% of the FVS perturbations when simulating steady states using SFA. This result is expected, as the SFA algorithm has an accuracy of about 60-80% for estimation of steady state values based only on topological information (Lee and Cho, 2018).

Additionally, the use of FC with source nodes and CIs produced by OCSANA improved upon the simulated results, increasing the correct simulation of the experimental results to 85%. The addition of graph theory source nodes to the optimal control set increases the power of the computational model to precisely simulate the activities in the actual embryo. Experimentally, *Gata.a* is necessary for zygotic development and ectodermal specification; although a source node in the GRN, *Gata.a* is activated in the ascidian embryo by animal sphere orientation (Satou and Imai, 2015). Thus, control of *Gata.a* in silico replicates a crucial signal in a biological system. We were unable to replicate the specification of the pan-neural cell fate using FVS perturbation adNze, or adNze with additional control of source nodes. This is mostly likely due to limitations of the GRN topology. In the GRN, only *Zic-r.b* activates pan-neural marker *Celf3a*. However, *Zic-r.b* is down-regulated in the adNze perturbation. The topological estimation of signal flow performed by SFA for *Celf3.a* is downregulated when compared with the unperturbed steady state because direct inhibition of *Zic-r.b* cannot be overcome by multiple activating signals into *Zic-r.b*. In Mochizuki et al. (2013), it is noted that if the FVS cannot explain all of the observed biological diversity in phenotypes, the issue may lie within the gene network structure. Furthermore, SFA is highly dependent on the network topology to correctly predict steady state activities of network nodes. Overall, the FC, SFA, and OCSANA algorithms in OCSANA+ can be used to simulate and prioritize *in vitro* perturbation experiments using solely the topological information provided by gene regulatory networks.

Finally, using the FC and SFA algorithms in OCSANA+, we are able to reproduce the results of FC of the Boolean simulation for the *Drosophila* segment polarity genes network in Zañudo et al. (2017) with the derived signaling network. We observe that by activating a set of FC nodes, we are able to upregulate all the correct patterning genes when compared to the unpatterned state. It is important to note that the ability of SFA to predict steady state values is highly dependent on the correct network topology. OCSANA+ provides a useful tool for simulating FC experiments when dynamic models may not be available.

